# A single-cell clusters similarity measure for different batches, datasets, and samples

**DOI:** 10.1101/2022.03.14.483731

**Authors:** Óscar González-Velasco, Elena Sanchez-Luis, Enrique De La Rosa, José Manuel Sánchez-Santos, Javier De Las Rivas

## Abstract

**Summary:** Since the inception of single-cell level measuring techniques, identification of distinct cell stages, phenotypes and populations has been a challenge. Cell clustering and dimensionality reduction methods are the most popular approaches to identify heterogeneity of single-cell data. But, as public repositories continue to grow in number, integrative analyses and merging of large pools of samples from different and heterogeneous datasets becomes a difficult challenge, which showcases the impossibility of scalability of some of the existing methods. Here we present *ClusterFoldSimilarity*, an R package that calculates a measure of similarity between clusters from different datasets/batches, without the need of correcting for batch effect or normalizing and merging the data, thus avoiding artifacts and the loss of information derived from these kinds of techniques. The similarity metric is based on the average vector module and sign of the product of logarithmic fold-changes. *ClusterFoldSimilarity* compares every single pair of clusters from any number of different samples/datasets, including different number of clusters for each sample. Additionally, the algorithm is able to select the top genes which contribute the most to the similarity of two specific clusters, serving also as a feature selection tool.

**Availability and implementation:** The algorithm is freely available as an R package at: https://github.com/OscarGVelasco/ClusterFoldSimilarity

## Introduction

Single-cell technologies (such as scRNA-seq, scATAC-seq or scCyToF) constitute a new collection of advanced methodologies capable of capturing the biomolecular state of the cell and measuring multiple cellular markers for hundreds to millions of cells. Using these technologies, researchers are able to elucidate genomic, epigenomic, and transcriptomic heterogeneity in cellular populations and subpopulations, and the variability at these levels (Kashima et al., 2020). The number of large-scale single-cell omic datasets generated using different technologies is continually growing with the advent of accessible datasets in public repositories. Despite this, integration of single-cell data remains difficult due to inherent dataset-specific systematic variability that present a challenge for batch-effect removal. A number of tools for batch-effect removal exist, but each has its advantages and limitations, with no clearly superior method (Tran et al., 2020). Here we present a new application tool that is able to analyze the similarity between clusters from different and heterogeneous datasets, without the need of batch-effect removal or merging. The approach is based on the fold change of the measured signal between clusters, standing on the idea that the overall signal change across different cell populations, subpopulations or phenotypes in a dataset should be conserved in different datasets measuring similar populations.

## Materials and methods

### Pairwise fold changes

In single-cell data, signal fold-changes (FC) can be defined as a Euclidean vector with origin at 0 and direction equal to the signal change, the scalar product of two vectors in the same space gives as a result a perpendicular vector to the plane. In the case of the dot product of two foldchanges the direction of the perpendicular vector will act as a concordance value of the direction of the FCs (see Figure 1A,B,C). The FCs are computed using the normalized count matrix. They are calculated pairwise between each cluster *i* and the other clusters belonging to the same sample. FC for each gene is calculated as the log2 of the ratio of the mean expression from cells in cluster *i* and the mean expression from cells in cluster *j*: ∀*j* ≠ *i*. In such manner, we obtain *k*-1 FCs (*k*: number of clusters in a given sample) for each of the clusters in that sample.

**Figure 1:**
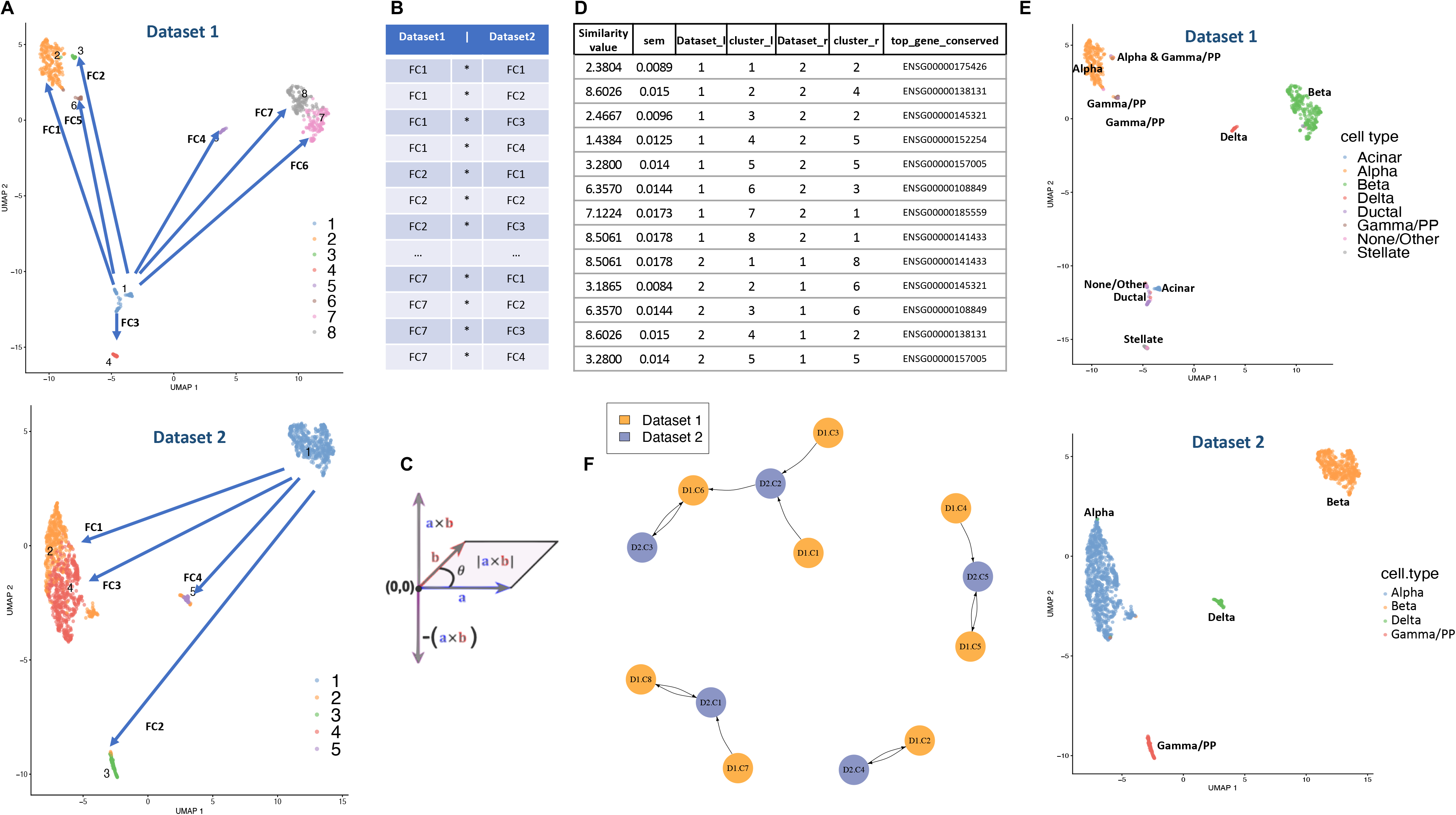
Pipeline representation of the similarity measurement done by ClusterFoldSimilarity. A: Foldchanges (FC) are calculated between all possible combination of clusters for each dataset, figure shows the FC calculation for cluster 1 in both datasets 1 and 2 in a UMAP corresponding with pancreas cells. B: The scalar product of the computed FCs between all pairs is calculated. C: Representation of the vectorial space of FCs and their product (in the graphic, a and b represent FCs of a same gene between clusters), FC that points in the same direction will have positive product sign, meanwhile opposite FC will have negative product sign. D: Table with the similarity values and output information given by ClusterFoldSimilarity. E: UMAP representation of two pancreatic datasets with annotated cell-types. Note that exact cell-type clusters are matched together by the table given at D. F: Graph plot made by ClusterFoldSimilarity using the similarity values calculated (given at table D), nodes correspond with clusters from a specific dataset, edges correspond with the similarity value, arrows point in the direction of the similarity relationship.

### Similarity measurement

Once all the possible combinations of FCs between all clusters from each sample/batch are computed, we start by comparing the cluster 1 from sample/batch 1 with the cluster 1 in sample/batch 2 and continue iteratively. We achieve this by computing the scalar product of every possible combination of FCs from the two clusters: *combinations* = (*k*_1_ − 1) × (*k*_2_ − 1) being *k*_1_: number of clusters in sample 1; *k*_2_: number of clusters in sample 2. Then we calculate for each gene the mean of the scalar products. This value will serve as a measure of gene importance, being the gene with the highest positive value the one that contributes the most to the similarity between the two clusters (negative values indicate opposite FC values).

Finally, the similarity value for a pair of clusters is calculated as the square root of the sum of the mean of genes (calculated in the previous step):

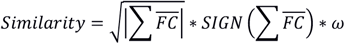

The weight *ω* is based on the concordance of positive versus negative FC’s and is calculated as:

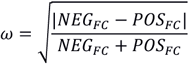

The *maximum similarity positive value* (calculated as described) will correspond with the most similar cluster: it is expected that, for a specific population of cells, overall changes in gene expression across different clusters (i.e., in different single cell datasets with similar cell populations) are conserved on different samples, and thus the sign of the product of FCs should be positive and its module higher than less concordant expression changes.

### Compatibility

*ClusterFoldSimilarity* is compatible with the most popular single-cell pipelines for an easy downstream integration: Seurat (Hao et al., 2021) and SingleCellExperiment (Amezquita et al., 2020).

## Results

*ClusterFoldSimilarity* has been tested in several single-cell datasets in which cell-type information was available. The first is comprised of four human pancreas datasets from the scRNAseq Bioconductor package (Risso D, 2021): GSE84133 (Baron et al., 2016); GSE86469 (Lawlor et al., 2017); GSE81608, (Xin et al., 2016); ArrayExpress: E-MTAB-5061 (Segerstolpe et al., 2016). Additionally, it has been tested in four peripheral blood mononuclear cell (PBMC) single-cell RNA-Seq datasets: GSE149938 (Xie et al., 2021); 10x Genomics 10k Peripheral blood mononuclear cells (PBMCs) from a healthy donor, Single Indexed; GSE120221 (Oetjen et al., 2018); PRJEB40448 (Lawlor et al., 2021).

Our results show that ClusterFoldSimilarity metric is able to pair clusters by cell type and cell subpopulation with high accuracy: all clusters corresponding with shared cell types across datasets are paired together, including subpopulations in the case of PBMCs. Moreover, the top gene markers selected by similarity correspond to markers reported on scientific literature (see Supp. Material).

## Supporting information

Supplementary material results

