## Supplementary material results for "A single-cell clusters similarity measure for different batches, datasets, and samples"

**Pancreatic Single-Cell datasets:**

A list of four human pancreatic single-cell RNA-Seq datasets from the scRNAseq package (Risso D, 2021) (GSE84133 (Baron et al., 2016), GSE86469 (Lawlor et al., 2017), GSE81608 (Xin et al., 2016), ArrayExpress: E-MTAB-5061 (Segerstolpe et al., 2016)) have been used to test ClusterFoldSimilarity performance. As shown in TABLE and FIGURE, clusters from different datasets corresponding with the same cell types get the highest similarity value among them (e.g., clusters 2 and 10 from dataset 1 GSE84133 (Baron et al., 2016) which correspond to alpha pancreatic cells both point to the cluster 2 in dataset 2, which corresponds as well with alpha pancreatic cells).

ClusterFoldSimilarity algorithm is also able to detect and match cell subpopulations across different datasets. Beta pancreatic cells are known to have different cell subpopulations (Dorrell et al.), these high specific subpopulations have a very similar profile expression. Clusters 4 and 6 of dataset 1 correspond with cluster 8 of dataset 2, meanwhile clusters 7 and 9 from dataset 1 correspond with cluster 7 of dataset 2 (see Supplementary Figure 1).

| Similarity value | sem | dataset_l | cluster_l | dataset_r | cluster_r | Top gene conserved |
|---|---|---|---|---|---|---|
| **8.1155** | 0.0539 | 1 | 1 | 2 | 4 | ENSG00000113140 |
| **1.3196** | 0.0204 | 1 | 1 | 3 | 5 | ENSG00000152254 |
| **7.6239** | 0.045 | 1 | 1 | 4 | 8 | ENSG00000113140 |
| **5.3361** | 0.0354 | 1 | 2 | 2 | 2 | ENSG00000115263 |
| **4.4394** | 0.0243 | 1 | 2 | 3 | 4 | ENSG00000170561 |
| **3.6455** | 0.0241 | 1 | 2 | 4 | 11 | ENSG00000170561 |
| **8.3194** | 0.0519 | 1 | 3 | 2 | 1 | ENSG00000148346 |
| **1.3410** | 0.022 | 1 | 3 | 3 | 5 | ENSG00000152254 |
| **7.6026** | 0.0427 | 1 | 3 | 4 | 2 | ENSG00000148346 |
| **4.4733** | 0.0273 | 1 | 4 | 2 | 8 | ENSG00000254647 |
| **3.7825** | 0.0297 | 1 | 4 | 3 | 1 | ENSG00000121351 |
| **4.2822** | 0.0266 | 1 | 4 | 4 | 1 | ENSG00000121351 |
| **8.5876** | 0.0638 | 1 | 5 | 2 | 1 | ENSG00000115386 |
| **1.2502** | 0.0249 | 1 | 5 | 3 | 5 | ENSG00000152254 |
| **8.2258** | 0.0624 | 1 | 5 | 4 | 6 | ENSG00000115386 |
| **4.0515** | 0.0288 | 1 | 6 | 2 | 8 | ENSG00000141433 |
| **3.8855** | 0.0322 | 1 | 6 | 3 | 1 | ENSG00000141433 |
| **3.3770** | 0.0278 | 1 | 6 | 4 | 1 | ENSG00000141433 |
| **5.2377** | 0.0344 | 1 | 7 | 2 | 7 | ENSG00000185559 |
| **4.0825** | 0.0344 | 1 | 7 | 3 | 1 | ENSG00000185559 |
| **4.5297** | 0.0316 | 1 | 7 | 4 | 1 | ENSG00000185559 |
| **5.1172** | 0.0397 | 1 | 8 | 2 | 5 | ENSG00000157005 |
| **3.1624** | 0.0347 | 1 | 8 | 3 | 5 | ENSG00000138207 |
| **1.9862** | 0.0159 | 1 | 8 | 4 | 10 | ENSG00000175426 |
| **5.4923** | 0.0322 | 1 | 9 | 2 | 7 | ENSG00000185559 |
| **4.2239** | 0.0354 | 1 | 9 | 3 | 1 | ENSG00000121351 |
| **4.4550** | 0.032 | 1 | 9 | 4 | 1 | ENSG00000121351 |
| **6.6343** | 0.0447 | 1 | 10 | 2 | 2 | ENSG00000169903 |
| **4.1278** | 0.0309 | 1 | 10 | 3 | 4 | ENSG00000138131 |
| **5.1411** | 0.0264 | 1 | 10 | 4 | 3 | ENSG00000163499 |
| **4.5525** | 0.052 | 1 | 11 | 2 | 6 | ENSG00000108849 |
| **3.6148** | 0.046 | 1 | 11 | 3 | 3 | ENSG00000108849 |
| **4.0686** | 0.0356 | 1 | 11 | 4 | 10 | ENSG00000108849 |
| **8.5876** | 0.0638 | 2 | 1 | 1 | 5 | ENSG00000115386 |
| **1.4933** | 0.0428 | 2 | 1 | 3 | 5 | ENSG00000124939 |
| **9.3213** | 0.0764 | 2 | 1 | 4 | 2 | ENSG00000148346 |
| **6.6343** | 0.0447 | 2 | 2 | 1 | 10 | ENSG00000169903 |
| **7.9140** | 0.055 | 2 | 2 | 3 | 4 | ENSG00000138131 |
| **8.3345** | 0.0562 | 2 | 2 | 4 | 11 | ENSG00000138131 |
| **4.2790** | 0.0288 | 2 | 3 | 1 | 2 | ENSG00000145321 |
| **4.4505** | 0.0414 | 2 | 3 | 3 | 4 | ENSG00000145321 |
| **3.6440** | 0.0367 | 2 | 3 | 4 | 9 | ENSG00000145321 |
| **8.1155** | 0.0539 | 2 | 4 | 1 | 1 | ENSG00000113140 |
| **1.6799** | 0.0474 | 2 | 4 | 3 | 5 | ENSG00000152254 |
| **12.3481** | 0.0954 | 2 | 4 | 4 | 8 | ENSG00000113140 |
| **5.1172** | 0.0397 | 2 | 5 | 1 | 8 | ENSG00000157005 |
| **4.2183** | 0.0557 | 2 | 5 | 3 | 5 | ENSG00000157005 |
| **3.7343** | 0.0523 | 2 | 5 | 4 | 1 | ENSG00000138207 |
| **4.5525** | 0.052 | 2 | 6 | 1 | 11 | ENSG00000108849 |

| | | | | | | |
|---|---|---|---|---|---|---|
| **4.3768** | 0.0575 | 2 | 6 | 3 | 3 | ENSG00000108849 |
| **5.8230** | 0.049 | 2 | 6 | 4 | 10 | ENSG00000108849 |
| **5.4923** | 0.0322 | 2 | 7 | 1 | 9 | ENSG00000185559 |
| **7.0179** | 0.064 | 2 | 7 | 3 | 1 | ENSG00000185559 |
| **7.9299** | 0.0581 | 2 | 7 | 4 | 1 | ENSG00000141433 |
| **5.3718** | 0.0334 | 2 | 8 | 1 | 9 | ENSG00000152254 |
| **7.4152** | 0.0644 | 2 | 8 | 3 | 1 | ENSG00000141433 |
| **9.4991** | 0.0611 | 2 | 8 | 4 | 1 | ENSG00000141433 |
| **4.2239** | 0.0354 | 3 | 1 | 1 | 9 | ENSG00000121351 |
| **7.4152** | 0.0644 | 3 | 1 | 2 | 8 | ENSG00000141433 |
| **6.6677** | 0.0619 | 3 | 1 | 4 | 1 | ENSG00000141433 |
| **3.9106** | 0.0271 | 3 | 2 | 1 | 10 | ENSG00000145321 |
| **5.0704** | 0.0463 | 3 | 2 | 2 | 2 | ENSG00000078098 |
| **3.0962** | 0.0267 | 3 | 2 | 4 | 3 | ENSG00000078098 |
| **3.6148** | 0.046 | 3 | 3 | 1 | 11 | ENSG00000108849 |
| **4.3768** | 0.0575 | 3 | 3 | 2 | 6 | ENSG00000108849 |
| **4.4488** | 0.0439 | 3 | 3 | 4 | 10 | ENSG00000108849 |
| **4.4394** | 0.0243 | 3 | 4 | 1 | 2 | ENSG00000170561 |
| **7.9140** | 0.055 | 3 | 4 | 2 | 2 | ENSG00000138131 |
| **6.7879** | 0.0414 | 3 | 4 | 4 | 11 | ENSG00000078098 |
| **3.1624** | 0.0347 | 3 | 5 | 1 | 8 | ENSG00000138207 |
| **4.2183** | 0.0557 | 3 | 5 | 2 | 5 | ENSG00000157005 |
| **2.4457** | 0.0408 | 3 | 5 | 4 | 2 | ENSG00000152254 |
| **4.5297** | 0.0316 | 4 | 1 | 1 | 7 | ENSG00000185559 |
| **9.4991** | 0.0611 | 4 | 1 | 2 | 8 | ENSG00000141433 |
| **6.6677** | 0.0619 | 4 | 1 | 3 | 1 | ENSG00000141433 |
| **7.6026** | 0.0427 | 4 | 2 | 1 | 3 | ENSG00000148346 |
| **9.3213** | 0.0764 | 4 | 2 | 2 | 1 | ENSG00000148346 |
| **2.4457** | 0.0408 | 4 | 2 | 3 | 5 | ENSG00000152254 |
| **5.1411** | 0.0264 | 4 | 3 | 1 | 10 | ENSG00000163499 |
| **6.9098** | 0.0429 | 4 | 3 | 2 | 2 | ENSG00000138131 |
| **4.5231** | 0.0312 | 4 | 3 | 3 | 4 | ENSG00000078098 |
| **4.3850** | 0.0313 | 4 | 4 | 1 | 10 | ENSG00000163499 |
| **7.7133** | 0.0514 | 4 | 4 | 2 | 2 | ENSG00000138131 |
| **5.6066** | 0.037 | 4 | 4 | 3 | 4 | ENSG00000138131 |
| **2.2650** | 0.0201 | 4 | 5 | 1 | 9 | ENSG00000121351 |
| **1.2880** | 0.0382 | 4 | 5 | 2 | 6 | ENSG00000108849 |
| **0.7766** | 0.0368 | 4 | 5 | 3 | 1 | ENSG00000121351 |
| **8.2258** | 0.0624 | 4 | 6 | 1 | 5 | ENSG00000115386 |
| **8.9845** | 0.0868 | 4 | 6 | 2 | 1 | ENSG00000148346 |
| **2.3369** | 0.0434 | 4 | 6 | 3 | 5 | ENSG00000152254 |
| **4.3378** | 0.0513 | 4 | 7 | 1 | 5 | ENSG00000115386 |
| **6.7860** | 0.0674 | 4 | 7 | 2 | 1 | ENSG00000148346 |
| **1.7763** | 0.0329 | 4 | 7 | 3 | 5 | ENSG00000004848 |
| **7.6239** | 0.045 | 4 | 8 | 1 | 1 | ENSG00000113140 |
| **12.3481** | 0.0954 | 4 | 8 | 2 | 4 | ENSG00000113140 |
| **2.4334** | 0.0635 | 4 | 8 | 3 | 1 | ENSG00000145321 |
| **4.1159** | 0.0295 | 4 | 9 | 1 | 10 | ENSG00000138131 |
| **7.4034** | 0.0491 | 4 | 9 | 2 | 2 | ENSG00000138131 |
| **6.0655** | 0.0357 | 4 | 9 | 3 | 4 | ENSG00000138131 |

| | | | | | | |
|---|---|---|---|---|---|---|
| **4.0686** | 0.0356 | 4 | 10 | 1 | 11 | ENSG00000108849 |
| **5.8230** | 0.049 | 4 | 10 | 2 | 6 | ENSG00000108849 |
| **4.4488** | 0.0439 | 4 | 10 | 3 | 3 | ENSG00000108849 |
| **4.4892** | 0.0317 | 4 | 11 | 1 | 10 | ENSG00000138131 |
| **8.3345** | 0.0562 | 4 | 11 | 2 | 2 | ENSG00000138131 |
| **6.7879** | 0.0414 | 4 | 11 | 3 | 4 | ENSG00000078098 |

*Supplementary Table 1: ClusterFoldSimilarity computed results of the comparison of the four pancreatic scRNA-Seq datasets.*

**PBMC's Single-Cell datasets:**

A list of three human PBMC (Peripheral blood mononuclear cells) single-cell RNA-Seq datasets have been used to test ClusterFoldSimilarity measurement (see Supplementary table 2). A high accuracy of matching cell types is achieved by the similarity measurement method presented in this paper with well-known populations like B lymphocytes which correspond to clusters 0, 4 and 3 in the different datasets, respectively, and with well-establish markers as CD20 or CD23 (Tabata et al., 2014). The same occurs with hematopoietic stem cells present in datasets 1 and 3 (clusters 2 and 9, respectively) and again with very well-known markers acting like anchors as CD34 or SMIM24 (Tonglin et al., 2022).

Even distinct high-specialized subpopulations are paired as similar like clusters 9, 0 and 1 from the 3 datasets, respectively, that correspond to classical monocytes (CD14, FCN1 and FPR1 markers as the best evaluated) or non-classical ones (clusters 10, 6 and 8) with highly specialized markers like FCGR3A or MS4A7) (Vierstra et al., 2020; Xue et al., 2022). Another example are the myeloid dendritic cells and plasmacytoid dendritic ones in Datasets 2 and 3. Clusters 8 and 13 from these two datasets, respectively correspond to myeloid dendritic ones, while clusters 10 and 14 are related to plasmacytoid ones. In this last case, markers like CLEC10A and FCER1A, or LILRA4 and CLEC4C are key genes to distinguish them (Villani et al., 2017) (see Supplementary table 3: measurement results, and Supplementary Figure 2, in which clusters and cell-types of each of the three datasets of PBMCs are depicted).

| DATASETS | AUTHORS / REFERENCES | DATA PUBLIC | REPOSITORY LOCATION | PLATFORM USED | NUMBER OF CELLS | NUMBER OF HEALTHY DONORS | TISSUE | SUPERVISED DATA (FACS) |
|---|---|---|---|---|---|---|---|---|
| **Dataset 1** | Xie X et al. Natl Sci Rev (2021) | 2020 | GEO (Gene Expression Omnibus): GSE149938 | Illumina HiSeq 4000 (Homo sapiens) | 7,643 | 21 | BM/ PBMC | Yes |
| **Dataset 2** | 10x Genomics (2020) | 2020 | 10x Genomics Datasets: "10k Peripheral blood mononuclear cells (PBMCs) from a healthy donor, Single Indexed" | Illumina NovaSeq 6000 (Homo sapiens) | 10,985 | 1 | PBMC | No |
| **Dataset 3** | Oetjen KA et al. JCI Insight (2018) | 2018 | GEO (Gene Expression Omnibus): GSE120221 | Illumina HiSeq 3000 (Homo sapiens) | 90,653 | 20 | BMMC | No |

*Supplementary Table 2: List of PBMCs scRNA-Seq datasets used for the testing of the similarity measure of ClusterFoldSimilarity.*

| similarity_value | sem | dataset_l | cluster_l | dataset_r | cluster_r | top_gene_conserved |
|---|---|---|---|---|---|---|
| **58.88666** | 0.029156 | 1 | 0 | 2 | 4 | MS4A1 |
| **58.88666** | 0.029156 | 1 | 0 | 2 | 4 | FCRL1 |
| **58.88666** | 0.029156 | 1 | 0 | 2 | 4 | FCER2 |
| **39.59577** | 0.012671 | 1 | 0 | 2 | 1 | CCR7 |
| **39.59577** | 0.012671 | 1 | 0 | 2 | 1 | FCER1G |
| **39.59577** | 0.012671 | 1 | 0 | 2 | 1 | CD300A |
| **57.87181** | 0.019807 | 1 | 0 | 3 | 3 | MS4A1 |
| **57.87181** | 0.019807 | 1 | 0 | 3 | 3 | CD79A |
| **57.87181** | 0.019807 | 1 | 0 | 3 | 3 | FCER2 |
| **40.29241** | 0.00945 | 1 | 0 | 3 | 4 | CCR7 |
| **40.29241** | 0.00945 | 1 | 0 | 3 | 4 | SPN |
| **40.29241** | 0.00945 | 1 | 0 | 3 | 4 | CD63 |
| **48.95695** | 0.015483 | 1 | 1 | 2 | 4 | CD79A |
| **48.95695** | 0.015483 | 1 | 1 | 2 | 4 | CD79B |
| **48.95695** | 0.015483 | 1 | 1 | 2 | 4 | CD22 |
| **47.72975** | 0.010232 | 1 | 1 | 2 | 1 | FCER1G |
| **47.72975** | 0.010232 | 1 | 1 | 2 | 1 | CD300A |
| **47.72975** | 0.010232 | 1 | 1 | 2 | 1 | MPEG1 |
| **49.90028** | 0.021415 | 1 | 1 | 3 | 11 | VPREB1 |
| **49.90028** | 0.021415 | 1 | 1 | 3 | 11 | IGLL1 |
| **49.90028** | 0.021415 | 1 | 1 | 3 | 11 | CD9 |
| **43.16096** | 0.011471 | 1 | 1 | 3 | 3 | CD79A |

| | | | | | | |
|---|---|---|---|---|---|---|
| **43.16096** | 0.011471 | 1 | 1 | 3 | 3 | CD79B |
| **43.16096** | 0.011471 | 1 | 1 | 3 | 3 | CD22 |
| **43.25624** | 0.012234 | 1 | 2 | 2 | 10 | FCER1A |
| **43.25624** | 0.012234 | 1 | 2 | 2 | 10 | KCNK17 |
| **43.25624** | 0.012234 | 1 | 2 | 2 | 10 | FLT3 |
| **36.33448** | 0.013047 | 1 | 2 | 2 | 8 | FCER1A |
| **36.33448** | 0.013047 | 1 | 2 | 2 | 8 | MGST1 |
| **36.33448** | 0.013047 | 1 | 2 | 2 | 8 | FLT3 |
| **56.79049** | 0.01704 | 1 | 2 | 3 | 9 | SMIM24 |
| **56.79049** | 0.01704 | 1 | 2 | 3 | 9 | CD34 |
| **56.79049** | 0.01704 | 1 | 2 | 3 | 9 | NPR3 |
| **52.17337** | 0.012994 | 1 | 2 | 3 | 16 | MGST1 |
| **52.17337** | 0.012994 | 1 | 2 | 3 | 16 | SMIM24 |
| **52.17337** | 0.012994 | 1 | 2 | 3 | 16 | TNFSF13B |
| **66.05114** | 0.022851 | 1 | 3 | 2 | 8 | CD33 |
| **66.05114** | 0.022851 | 1 | 3 | 2 | 8 | FCN1 |
| **66.05114** | 0.022851 | 1 | 3 | 2 | 8 | IL13RA1 |
| **64.70186** | 0.027343 | 1 | 3 | 2 | 0 | FCN1 |
| **64.70186** | 0.027343 | 1 | 3 | 2 | 0 | CD14 |
| **64.70186** | 0.027343 | 1 | 3 | 2 | 0 | MS4A6A |
| **61.54175** | 0.018633 | 1 | 3 | 3 | 13 | FCN1 |
| **61.54175** | 0.018633 | 1 | 3 | 3 | 13 | FCER1G |
| **61.54175** | 0.018633 | 1 | 3 | 3 | 13 | CSF2RA |
| **58.23276** | 0.022307 | 1 | 3 | 3 | 1 | FCN1 |
| **58.23276** | 0.022307 | 1 | 3 | 3 | 1 | CD14 |
| **58.23276** | 0.022307 | 1 | 3 | 3 | 1 | MS4A6A |
| **27.2258** | 0.01144 | 1 | 4 | 2 | 10 | FLT3 |
| **27.2258** | 0.01144 | 1 | 4 | 2 | 10 | KCNK17 |
| **27.2258** | 0.01144 | 1 | 4 | 2 | 10 | RNF130 |
| **26.45601** | 0.011758 | 1 | 4 | 2 | 8 | CSF3R |
| **26.45601** | 0.011758 | 1 | 4 | 2 | 8 | FLT3 |
| **26.45601** | 0.011758 | 1 | 4 | 2 | 8 | RNF130 |
| **55.12599** | 0.014773 | 1 | 4 | 3 | 9 | SMIM24 |
| **55.12599** | 0.014773 | 1 | 4 | 3 | 9 | CD34 |
| **55.12599** | 0.014773 | 1 | 4 | 3 | 9 | NPR3 |
| **47.46057** | 0.010639 | 1 | 4 | 3 | 16 | SMIM24 |
| **47.46057** | 0.010639 | 1 | 4 | 3 | 16 | CSF3R |
| **47.46057** | 0.010639 | 1 | 4 | 3 | 16 | MGST1 |
| **54.77731** | 0.020701 | 1 | 5 | 2 | 4 | CD79A |
| **54.77731** | 0.020701 | 1 | 5 | 2 | 4 | MS4A1 |
| **54.77731** | 0.020701 | 1 | 5 | 2 | 4 | FCRL1 |
| **38.28739** | 0.00986 | 1 | 5 | 2 | 1 | FCER1G |
| **38.28739** | 0.00986 | 1 | 5 | 2 | 1 | CD300A |
| **38.28739** | 0.00986 | 1 | 5 | 2 | 1 | RNF130 |
| **51.3691** | 0.014009 | 1 | 5 | 3 | 3 | CD79A |

| | | | | | | |
|---|---|---|---|---|---|---|
| **51.3691** | 0.014009 | 1 | 5 | 3 | 3 | MS4A1 |
| **51.3691** | 0.014009 | 1 | 5 | 3 | 3 | CD79B |
| **51.16881** | 0.021266 | 1 | 5 | 3 | 11 | VPREB1 |
| **51.16881** | 0.021266 | 1 | 5 | 3 | 11 | CD9 |
| **51.16881** | 0.021266 | 1 | 5 | 3 | 11 | MME |
| **42.1961** | 0.015063 | 1 | 6 | 2 | 10 | FCER1A |
| **42.1961** | 0.015063 | 1 | 6 | 2 | 10 | CSF2RB |
| **42.1961** | 0.015063 | 1 | 6 | 2 | 10 | RNF130 |
| **33.75787** | 0.015632 | 1 | 6 | 2 | 8 | FCER1A |
| **33.75787** | 0.015632 | 1 | 6 | 2 | 8 | RNF130 |
| **33.75787** | 0.015632 | 1 | 6 | 2 | 8 | SLC40A1 |
| **58.1692** | 0.015619 | 1 | 6 | 3 | 9 | NPR3 |
| **58.1692** | 0.015619 | 1 | 6 | 3 | 9 | SLC40A1 |
| **58.1692** | 0.015619 | 1 | 6 | 3 | 9 | SLC39A3 |
| **54.62244** | 0.012812 | 1 | 6 | 3 | 16 | MGST1 |
| **54.62244** | 0.012812 | 1 | 6 | 3 | 16 | EREG |
| **54.62244** | 0.012812 | 1 | 6 | 3 | 16 | RNF130 |
| **59.02517** | 0.022631 | 1 | 7 | 2 | 1 | CD3D |
| **59.02517** | 0.022631 | 1 | 7 | 2 | 1 | MAL |
| **59.02517** | 0.022631 | 1 | 7 | 2 | 1 | IL7R |
| **58.90577** | 0.020905 | 1 | 7 | 2 | 3 | CD3D |
| **58.90577** | 0.020905 | 1 | 7 | 2 | 3 | IL7R |
| **58.90577** | 0.020905 | 1 | 7 | 2 | 3 | CD3G |
| **59.11541** | 0.01632 | 1 | 7 | 3 | 10 | CD3D |
| **59.11541** | 0.01632 | 1 | 7 | 3 | 10 | CD3E |
| **59.11541** | 0.01632 | 1 | 7 | 3 | 10 | CD3G |
| **58.71306** | 0.017002 | 1 | 7 | 3 | 0 | CD3D |
| **58.71306** | 0.017002 | 1 | 7 | 3 | 0 | CD3E |
| **58.71306** | 0.017002 | 1 | 7 | 3 | 0 | IL7R |
| **65.61133** | 0.037952 | 1 | 9 | 2 | 0 | CD14 |
| **65.61133** | 0.037952 | 1 | 9 | 2 | 0 | FCN1 |
| **65.61133** | 0.037952 | 1 | 9 | 2 | 0 | FPR1 |
| **63.53123** | 0.027933 | 1 | 9 | 2 | 8 | FCN1 |
| **63.53123** | 0.027933 | 1 | 9 | 2 | 8 | CLEC7A |
| **63.53123** | 0.027933 | 1 | 9 | 2 | 8 | MS4A6A |
| **60.86746** | 0.030671 | 1 | 9 | 3 | 1 | FCN1 |
| **60.86746** | 0.030671 | 1 | 9 | 3 | 1 | CD14 |
| **60.86746** | 0.030671 | 1 | 9 | 3 | 1 | CLEC7A |
| **60.76157** | 0.020919 | 1 | 9 | 3 | 13 | FCN1 |
| **60.76157** | 0.020919 | 1 | 9 | 3 | 13 | FCER1G |
| **60.76157** | 0.020919 | 1 | 9 | 3 | 13 | CSF2RA |
| **64.28206** | 0.033541 | 1 | 10 | 2 | 0 | FCN1 |
| **64.28206** | 0.033541 | 1 | 10 | 2 | 0 | CLEC7A |
| **64.28206** | 0.033541 | 1 | 10 | 2 | 0 | C5AR1 |
| **62.01679** | 0.037604 | 1 | 10 | 2 | 6 | FCGR3A |

| | | | | | | |
|---|---|---|---|---|---|---|
| **62.01679** | 0.037604 | 1 | 10 | 2 | 6 | MS4A7 |
| **62.01679** | 0.037604 | 1 | 10 | 2 | 6 | CSF1R |
| **59.55731** | 0.026531 | 1 | 10 | 3 | 1 | FCN1 |
| **59.55731** | 0.026531 | 1 | 10 | 3 | 1 | CLEC7A |
| **59.55731** | 0.026531 | 1 | 10 | 3 | 1 | FCER1G |
| **57.86065** | 0.036922 | 1 | 10 | 3 | 8 | FCGR3A |
| **57.86065** | 0.036922 | 1 | 10 | 3 | 8 | MS4A7 |
| **57.86065** | 0.036922 | 1 | 10 | 3 | 8 | LILRB2 |
| **52.6742** | 0.015921 | 1 | 11 | 2 | 8 | MGST1 |
| **52.6742** | 0.015921 | 1 | 11 | 2 | 8 | CSF3R |
| **52.6742** | 0.015921 | 1 | 11 | 2 | 8 | CD33 |
| **49.45413** | 0.014888 | 1 | 11 | 2 | 10 | IL3RA |
| **49.45413** | 0.014888 | 1 | 11 | 2 | 10 | ALCAM |
| **49.45413** | 0.014888 | 1 | 11 | 2 | 10 | FLT3 |
| **56.40312** | 0.017683 | 1 | 11 | 3 | 9 | SMIM24 |
| **56.40312** | 0.017683 | 1 | 11 | 3 | 9 | MGST1 |
| **56.40312** | 0.017683 | 1 | 11 | 3 | 9 | CSF3R |
| **55.54274** | 0.01998 | 1 | 11 | 3 | 16 | MS4A3 |
| **55.54274** | 0.01998 | 1 | 11 | 3 | 16 | MGST1 |
| **55.54274** | 0.01998 | 1 | 11 | 3 | 16 | SMIM24 |
| **39.8598** | 0.01028 | 1 | 13 | 2 | 2 | IL7R |
| **39.8598** | 0.01028 | 1 | 13 | 2 | 2 | TNFRSF1B |
| **39.8598** | 0.01028 | 1 | 13 | 2 | 2 | MPEG1 |
| **39.43784** | 0.009852 | 1 | 13 | 2 | 1 | IL7R |
| **39.43784** | 0.009852 | 1 | 13 | 2 | 1 | FCER1G |
| **39.43784** | 0.009852 | 1 | 13 | 2 | 1 | MPEG1 |
| **49.76142** | 0.010637 | 1 | 13 | 3 | 9 | SMIM24 |
| **49.76142** | 0.010637 | 1 | 13 | 3 | 9 | IGLL1 |
| **49.76142** | 0.010637 | 1 | 13 | 3 | 9 | FLT3 |
| **46.67279** | 0.015733 | 1 | 13 | 3 | 11 | VPREB1 |
| **46.67279** | 0.015733 | 1 | 13 | 3 | 11 | IGLL1 |
| **46.67279** | 0.015733 | 1 | 13 | 3 | 11 | CD79A |
| **57.50837** | 0.025433 | 1 | 14 | 2 | 5 | CCL5 |
| **57.50837** | 0.025433 | 1 | 14 | 2 | 5 | CD8B |
| **57.50837** | 0.025433 | 1 | 14 | 2 | 5 | CD8A |
| **53.46948** | 0.018527 | 1 | 14 | 2 | 3 | CD3D |
| **53.46948** | 0.018527 | 1 | 14 | 2 | 3 | CD3G |
| **53.46948** | 0.018527 | 1 | 14 | 2 | 3 | CD3E |
| **55.8944** | 0.016602 | 1 | 14 | 3 | 7 | CCL5 |
| **55.8944** | 0.016602 | 1 | 14 | 3 | 7 | KLRD1 |
| **55.8944** | 0.016602 | 1 | 14 | 3 | 7 | CD3E |
| **55.68787** | 0.019206 | 1 | 14 | 3 | 2 | CCL5 |
| **55.68787** | 0.019206 | 1 | 14 | 3 | 2 | CD8A |
| **55.68787** | 0.019206 | 1 | 14 | 3 | 2 | CD8B |
| **58.22855** | 0.028945 | 1 | 15 | 2 | 4 | MS4A1 |

| | | | | | | |
|---|---|---|---|---|---|---|
| **58.22855** | 0.028945 | 1 | 15 | 2 | 4 | TNFRSF13C |
| **58.22855** | 0.028945 | 1 | 15 | 2 | 4 | CD79A |
| **29.67392** | 0.01347 | 1 | 15 | 2 | 1 | FCER1G |
| **29.67392** | 0.01347 | 1 | 15 | 2 | 1 | CD27 |
| **29.67392** | 0.01347 | 1 | 15 | 2 | 1 | CCR7 |
| **57.27898** | 0.01981 | 1 | 15 | 3 | 3 | MS4A1 |
| **57.27898** | 0.01981 | 1 | 15 | 3 | 3 | CD79A |
| **57.27898** | 0.01981 | 1 | 15 | 3 | 3 | TNFRSF13C |
| **39.03069** | 0.009558 | 1 | 15 | 3 | 10 | IL2RA |
| **39.03069** | 0.009558 | 1 | 15 | 3 | 10 | CD27 |
| **39.03069** | 0.009558 | 1 | 15 | 3 | 10 | RNF130 |
| **50.5379** | 0.021051 | 1 | 16 | 2 | 4 | CD79A |
| **50.5379** | 0.021051 | 1 | 16 | 2 | 4 | TNFRSF13B |
| **50.5379** | 0.021051 | 1 | 16 | 2 | 4 | MS4A1 |
| **24.92892** | 0.01349 | 1 | 16 | 2 | 3 | CD27 |
| **24.92892** | 0.01349 | 1 | 16 | 2 | 3 | FCER1G |
| **24.92892** | 0.01349 | 1 | 16 | 2 | 3 | SLAMF1 |
| **49.70222** | 0.032415 | 1 | 16 | 3 | 15 | DERL3 |
| **49.70222** | 0.032415 | 1 | 16 | 3 | 15 | TNFRSF17 |
| **49.70222** | 0.032415 | 1 | 16 | 3 | 15 | SDC1 |
| **45.25872** | 0.013778 | 1 | 16 | 3 | 3 | CD79A |
| **45.25872** | 0.013778 | 1 | 16 | 3 | 3 | MS4A1 |
| **45.25872** | 0.013778 | 1 | 16 | 3 | 3 | TNFRSF13B |
| **55.11914** | 0.030916 | 1 | 17 | 2 | 7 | KLRD1 |
| **55.11914** | 0.030916 | 1 | 17 | 2 | 7 | KLRF1 |
| **55.11914** | 0.030916 | 1 | 17 | 2 | 7 | CCL5 |
| **53.30639** | 0.021536 | 1 | 17 | 2 | 5 | CCL5 |
| **53.30639** | 0.021536 | 1 | 17 | 2 | 5 | KLRD1 |
| **53.30639** | 0.021536 | 1 | 17 | 2 | 5 | CD247 |
| **53.87584** | 0.02319 | 1 | 17 | 3 | 7 | KLRD1 |
| **53.87584** | 0.02319 | 1 | 17 | 3 | 7 | CCL5 |
| **53.87584** | 0.02319 | 1 | 17 | 3 | 7 | KLRF1 |
| **53.25009** | 0.024402 | 1 | 17 | 3 | 5 | KLRD1 |
| **53.25009** | 0.024402 | 1 | 17 | 3 | 5 | KLRF1 |
| **53.25009** | 0.024402 | 1 | 17 | 3 | 5 | CCL5 |
| **62.85813** | 0.029285 | 2 | 0 | 3 | 1 | CD14 |
| **62.85813** | 0.029285 | 2 | 0 | 3 | 1 | FCN1 |
| **62.85813** | 0.029285 | 2 | 0 | 3 | 1 | MS4A6A |
| **61.11947** | 0.020852 | 2 | 0 | 3 | 13 | IFI30 |
| **61.11947** | 0.020852 | 2 | 0 | 3 | 13 | FCN1 |
| **61.11947** | 0.020852 | 2 | 0 | 3 | 13 | MS4A6A |
| **62.56354** | 0.013077 | 2 | 1 | 3 | 0 | IL7R |
| **62.56354** | 0.013077 | 2 | 1 | 3 | 0 | CCR7 |
| **62.56354** | 0.013077 | 2 | 1 | 3 | 0 | MAL |
| **59.61305** | 0.012409 | 2 | 1 | 3 | 10 | MAL |

| | | | | | | |
|---|---|---|---|---|---|---|
| **59.61305** | 0.012409 | 2 | 1 | 3 | 10 | IL7R |
| **59.61305** | 0.012409 | 2 | 1 | 3 | 10 | CD3D |
| **57.99968** | 0.0127 | 2 | 2 | 3 | 0 | IL7R |
| **57.99968** | 0.0127 | 2 | 2 | 3 | 0 | CCR7 |
| **57.99968** | 0.0127 | 2 | 2 | 3 | 0 | CD3D |
| **56.05179** | 0.019102 | 2 | 2 | 3 | 4 | CD8B |
| **56.05179** | 0.019102 | 2 | 2 | 3 | 4 | CD8A |
| **56.05179** | 0.019102 | 2 | 2 | 3 | 4 | CCR7 |
| **61.04866** | 0.012536 | 2 | 3 | 3 | 10 | IL7R |
| **61.04866** | 0.012536 | 2 | 3 | 3 | 10 | CD3D |
| **61.04866** | 0.012536 | 2 | 3 | 3 | 10 | CD3G |
| **59.64372** | 0.013675 | 2 | 3 | 3 | 6 | IL7R |
| **59.64372** | 0.013675 | 2 | 3 | 3 | 6 | TNFRSF4 |
| **59.64372** | 0.013675 | 2 | 3 | 3 | 6 | CD3D |
| **55.82491** | 0.024558 | 2 | 4 | 3 | 3 | MS4A1 |
| **55.82491** | 0.024558 | 2 | 4 | 3 | 3 | CD79A |
| **55.82491** | 0.024558 | 2 | 4 | 3 | 3 | TNFRSF13C |
| **47.36664** | 0.019311 | 2 | 4 | 3 | 11 | CD79A |
| **47.36664** | 0.019311 | 2 | 4 | 3 | 11 | CD79B |
| **47.36664** | 0.019311 | 2 | 4 | 3 | 11 | CD19 |
| **55.0916** | 0.015613 | 2 | 5 | 3 | 2 | CCL5 |
| **55.0916** | 0.015613 | 2 | 5 | 3 | 2 | CD8A |
| **55.0916** | 0.015613 | 2 | 5 | 3 | 2 | CD8B |
| **55.09063** | 0.015287 | 2 | 5 | 3 | 7 | CCL5 |
| **55.09063** | 0.015287 | 2 | 5 | 3 | 7 | KLRD1 |
| **55.09063** | 0.015287 | 2 | 5 | 3 | 7 | FCGR3A |
| **59.98892** | 0.033535 | 2 | 6 | 3 | 8 | MS4A7 |
| **59.98892** | 0.033535 | 2 | 6 | 3 | 8 | FCGR3A |
| **59.98892** | 0.033535 | 2 | 6 | 3 | 8 | C5AR1 |
| **59.41426** | 0.023695 | 2 | 6 | 3 | 1 | CLEC7A |
| **59.41426** | 0.023695 | 2 | 6 | 3 | 1 | FCN1 |
| **59.41426** | 0.023695 | 2 | 6 | 3 | 1 | C5AR1 |
| **53.08108** | 0.021444 | 2 | 7 | 3 | 7 | KLRD1 |
| **53.08108** | 0.021444 | 2 | 7 | 3 | 7 | KLRF1 |
| **53.08108** | 0.021444 | 2 | 7 | 3 | 7 | CCL5 |
| **52.62745** | 0.023964 | 2 | 7 | 3 | 5 | KLRD1 |
| **52.62745** | 0.023964 | 2 | 7 | 3 | 5 | KLRF1 |
| **52.62745** | 0.023964 | 2 | 7 | 3 | 5 | KLRB1 |
| **61.08256** | 0.023754 | 2 | 8 | 3 | 13 | CLEC10A |
| **61.08256** | 0.023754 | 2 | 8 | 3 | 13 | FCER1A |
| **61.08256** | 0.023754 | 2 | 8 | 3 | 13 | CD1C |
| **60.98436** | 0.02096 | 2 | 8 | 3 | 1 | FCN1 |
| **60.98436** | 0.02096 | 2 | 8 | 3 | 1 | CLEC7A |
| **60.98436** | 0.02096 | 2 | 8 | 3 | 1 | MS4A6A |
| **52.53756** | 0.030663 | 2 | 10 | 3 | 14 | LILRA4 |

| | | | | | | |
|---|---|---|---|---|---|---|
| **52.53756** | 0.030663 | 2 | 10 | 3 | 14 | CLEC4C |
| **52.53756** | 0.030663 | 2 | 10 | 3 | 14 | IL3RA |
| **50.4783** | 0.012019 | 2 | 10 | 3 | 16 | TNFSF13B |
| **50.4783** | 0.012019 | 2 | 10 | 3 | 16 | MS4A6A |
| **50.4783** | 0.012019 | 2 | 10 | 3 | 16 | RNF130 |

*Supplementary Table 3: ClusterFoldSimilarity computed results of the comparison of the three PBMCs scRNA-Seq datasets listed on supplementary table 2.*
