## Supplementary figures and images for "A single-cell clusters similarity measure for different batches, datasets, and samples"

### Supp. Figure S1

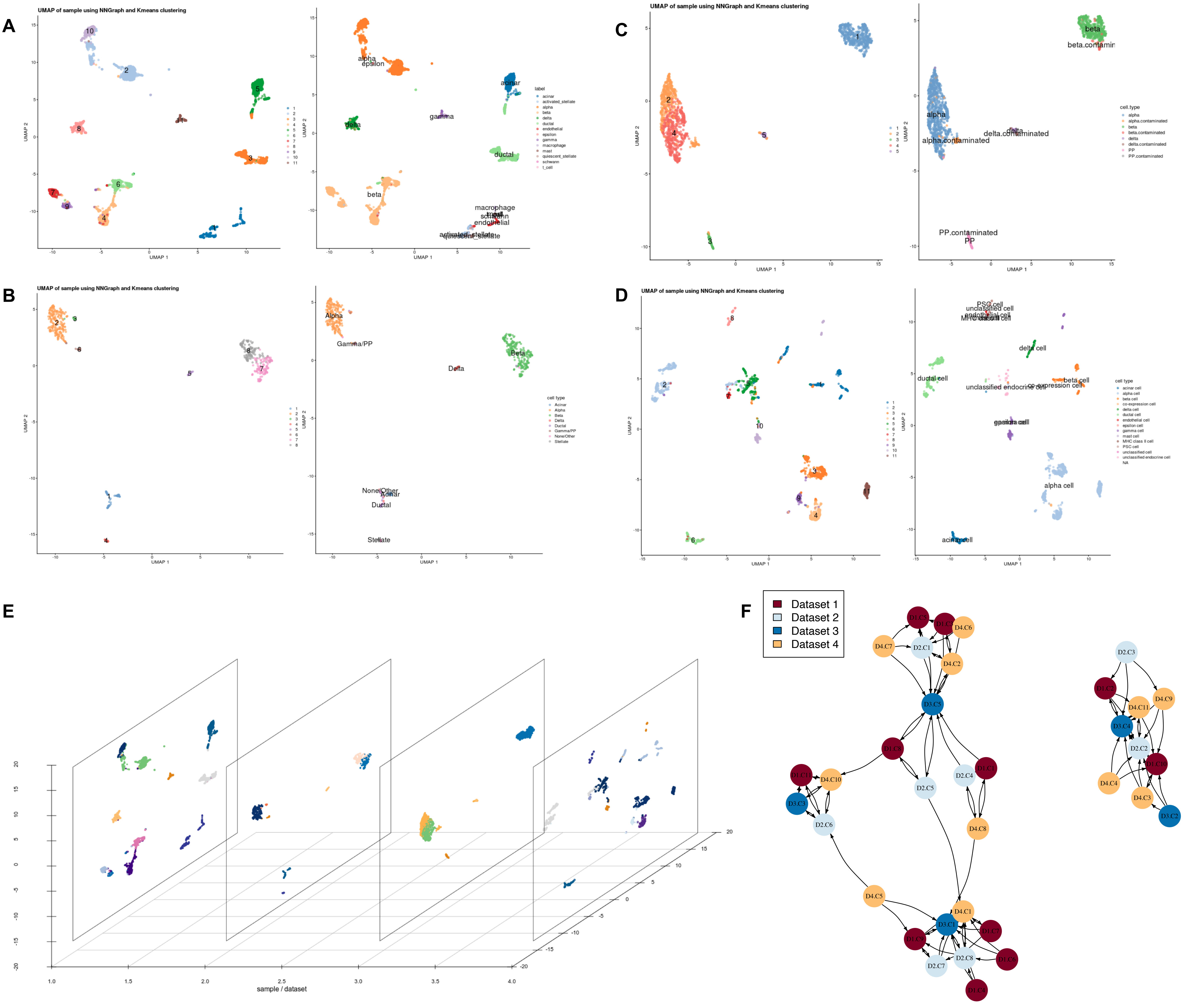

### Supp. Figure S2

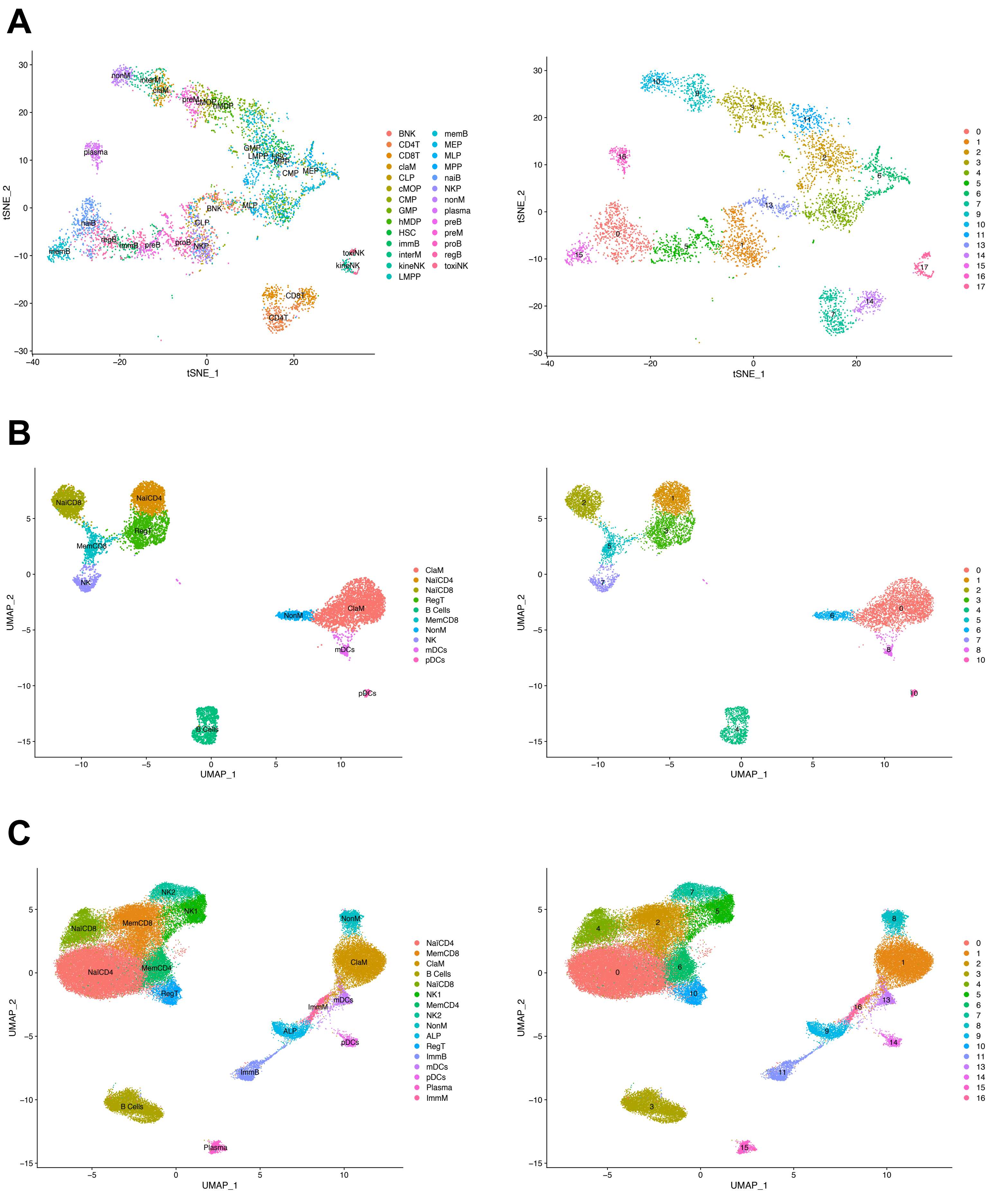
